# RNF152 Negatively Regulates mTOR Signalling and Blocks Cell Proliferation in the Floor Plate

**DOI:** 10.1101/646661

**Authors:** Minori Kadoya, Noriaki Sasai

## Abstract

The neural tube is composed of a number of neural progenitors and postmitotic neurons distributed in a quantitatively and spatially precise manner. The floor plate, located in the ventral-most region of the neural tube, has a lot of unique characteristics, including a low cell proliferation rate. The mechanisms by which this region-specific proliferation rate is regulated remain elusive.

Here we show that the activity of the mTOR signalling pathway, which regulates the proliferation of the neural progenitor cells, is significantly lower in the floor plate than in other domains of the embryonic neural tube. We identified the forkhead-type transcription factor FoxA2 as a negative regulator of mTOR signalling in the floor plate. We demonstrate that FoxA2 transcriptionally induces the expression of the E3 ubiquitin ligase RNF152, which together with its substrate RagA, regulates cell proliferation via the mTOR pathway. Silencing of RNF152 led to the aberrant upregulation of the mTOR signal and aberrant cell division in the floor plate. Taken together, the present findings suggest that floor plate cell number is controlled by the negative regulation of mTOR signalling through the activity of FoxA2 and its downstream effector RNF152.

## Introduction

The neural tube is the embryonic precursor to the central nervous system, and is composed of neural progenitor cells and postmitotic neurons (Le Dreau and Marti, 2012; Ribes and Briscoe, 2009). These cells are arranged in a quantitatively and spatially precise manner, which ensures that the neural tube develops into a functional organ.

During neural tube development, cell specification, proliferation and tissue growth are coordinated by secreted factors, collectively called morphogens (Dessaud et al., 2008; Kicheva et al., 2014; Perrimon et al., 2012). Among them, Sonic Hedgehog (Shh) is expressed in the floor plate (FP) and its underlying mesodermal tissue notochord, and the protein is distributed in a gradient from the ventral to the dorsal regions, with the highest level in the FP (Ribes and Briscoe, 2009). Each ventral progenitor cell acquires its own identity depending on the concentration of Shh (Dessaud et al., 2008; Jacob and Briscoe, 2003).

Shh regulates cell proliferation in parallel with the cell specification (Komada, 2012). Embryos devoid of the *Shh* gene exhibit not only defective pattern formation but also the reduced size of the neural tube, suggesting that Shh plays indispensable roles in both cell proliferation and tissue growth (Bulgakov et al., 2004; Chiang et al., 1996). However, sustained and excessive Shh signalling lead to tumorigenesis (Dahmane et al., 2001; Rowitch et al., 1999). The Shh signal, therefore, needs to be strictly regulated both spatially and temporally.

The FP, located at the ventral-most part of the neural tube, is a source of Shh, and acts as an organiser for the dorsal-ventral pattern formation of the neural tube (Dessaud et al., 2010; Yu et al., 2013). In addition, the FP has a number of unique characteristics compared with other neural domains (Placzek and Briscoe, 2005). At the trunk level, the FP is non-neurogenic (Ono et al., 2007), which is distinct from other progenitor domains where these cells differentiate into the corresponding neurons (Dessaud et al., 2008; Ribes and Briscoe, 2009). FP cells express guidance molecules such as Netrin and DCC, and which are essential for the precise guidance of the commissural axons (de la Torre et al., 1997; Kennedy et al., 1994; Ming et al., 1997; Sloan et al., 2015). The FP also expresses the actin-related factors, and is important for defining the neural tube shape (Nishimura et al., 2012; Nishimura and Takeichi, 2008). Together, the FP is indispensable for pattern formation, morphology, and functional control of the entire neural tube.

Neural progenitor cells in any neural domain actively proliferate and dynamically increases in number, whereas the FP cells, which are exposed to the highest level of Shh, do not actively increase (Kicheva et al., 2014). One possible explanation for this phenomenon is that the presence of a negative regulator(s) for the cell proliferation that is exclusively expressed in the FP region, and antagonise the proliferative effect of Shh.

The mechanistic target of rapamycin (mTOR) pathway is a versatile signalling system involved in a number of biological events including cell proliferation, survival and metabolism through early embryonic to postnatal stages (Gangloff et al., 2004; Laplante and Sabatini, 2009; Murakami et al., 2004). The mTORC (mTOR complex) is the hub of mTOR signal (Laplante and Sabatini, 2012), and acts as a serine/threonine kinase. Unsurprisingly, mTOR signal is essential for proper development of the central nervous system (LiCausi and Hartman, 2018; Ryskalin et al., 2017; Tee et al., 2016), and aberrant mTOR signalling is associated with neural defects during development. Blocking the mTOR signal with the phosphoinositide 3-kinase and mTOR inhibitors represses neurogenesis (Fishwick et al., 2010). Genetic elimination of the mTOR signal disrupts progenitor self-renewal and brain morphogenesis (Ka et al., 2014). Analysis of Tuberous sclerosis complex subunit 1 (Tsc1), a negative regulator of mTOR signaling (Dalle Pezze et al., 2012), shows that *Tsc1* homozygous mutant mice exhibit embryonic lethality with an unclosed neural tube (Kobayashi et al., 2001; Rennebeck et al., 1998).

In the neural tube, the mTOR pathway is active in ventral regions and in migrating neural crest cells, as shown by the expression of the phosphorylated form of mTOR (Nie et al., 2018). Because Shh is important for the assignment of ventral neural domains (Ribes and Briscoe, 2009) and migration of the neural crest (Kahane et al., 2013), the distribution of activated mTOR suggests an association between Shh and the mTOR signaling pathway.

The mTOR pathway phosphorylates and activates the transcription factor Gli1, a mediator of intracellular Shh signaling, and promotes the expression of target genes related to cell proliferation (Wang et al., 2012), thus supporting the relationship between mTOR and Shh signalling. Gli1 activation by mTOR is recognised as non-canonical in terms of the Gli activation, as this pathway is independent from the one mediated by the receptor protein for the Shh signal, Smoothened (Smo) (Dessaud et al., 2008). However, this signal pathway was demonstrated at the cellular level, and whether this pathway is also functional in a developmental context remains elusive. Abrogation of cilia activates the mTOR signal (Foerster et al., 2017), and Shh signaling requires cilia (Sasai and Briscoe, 2012), suggesting that Shh and mTOR have reciprocal activities. However, experiments in conditional knockout mice suggest that the underlying regulatory mechanism is context-dependent (Foerster et al., 2017).

In the present study, we mainly used chick embryos to investigate the mechanisms underlying the selective low proliferation rate of FP cells, with particular focus on the relationship between the Shh and mTOR signalling pathways. FoxA2, a transcription factor expressed in the FP and a target gene of Shh, blocked the mTOR signal, thereby altering cell proliferation. We identified the E3 ubiquitin ligase RNF152 as a target gene of FoxA2, and showed that RNF152 negatively regulates mTOR signalling by catalyzing the ubiquitination of the small GTPase RagA. Loss-of-function experiments were performed to demonstrate the role of RNF152 in regulating the proliferation of FP cells.

## Results

### The FP is significantly less proliferative than other neural domains

To clarify the mechanisms underlying the regulation of cell proliferation and tissue growth of the neural tube, the distribution of mitotic cells was examined by immunohistochemical detection of phospho-Histone 3 (Ser 10) (pHH3)-positive cells in cross sections of the neural tube. Embryos were harvested at Hamburger and Hamilton (HH) stage 11, soon after neural tube closure, HH stage 16, at the start of neurogenesis, and HH stage 22, when the neural tube matures; and pHH3 expression was analyzed at the anterior thoracic level. pHH3-positive cells were detected among apical cells along the dorsal-ventral axis (Figures 1A,B,C). However, pHH3-positive cells were not detected in the FP domain, which is characterized by high FoxA2 expression, at any stage (Figure 1A’,B’,C’), suggesting that FP cells were not proliferative.

**Figure 1.**
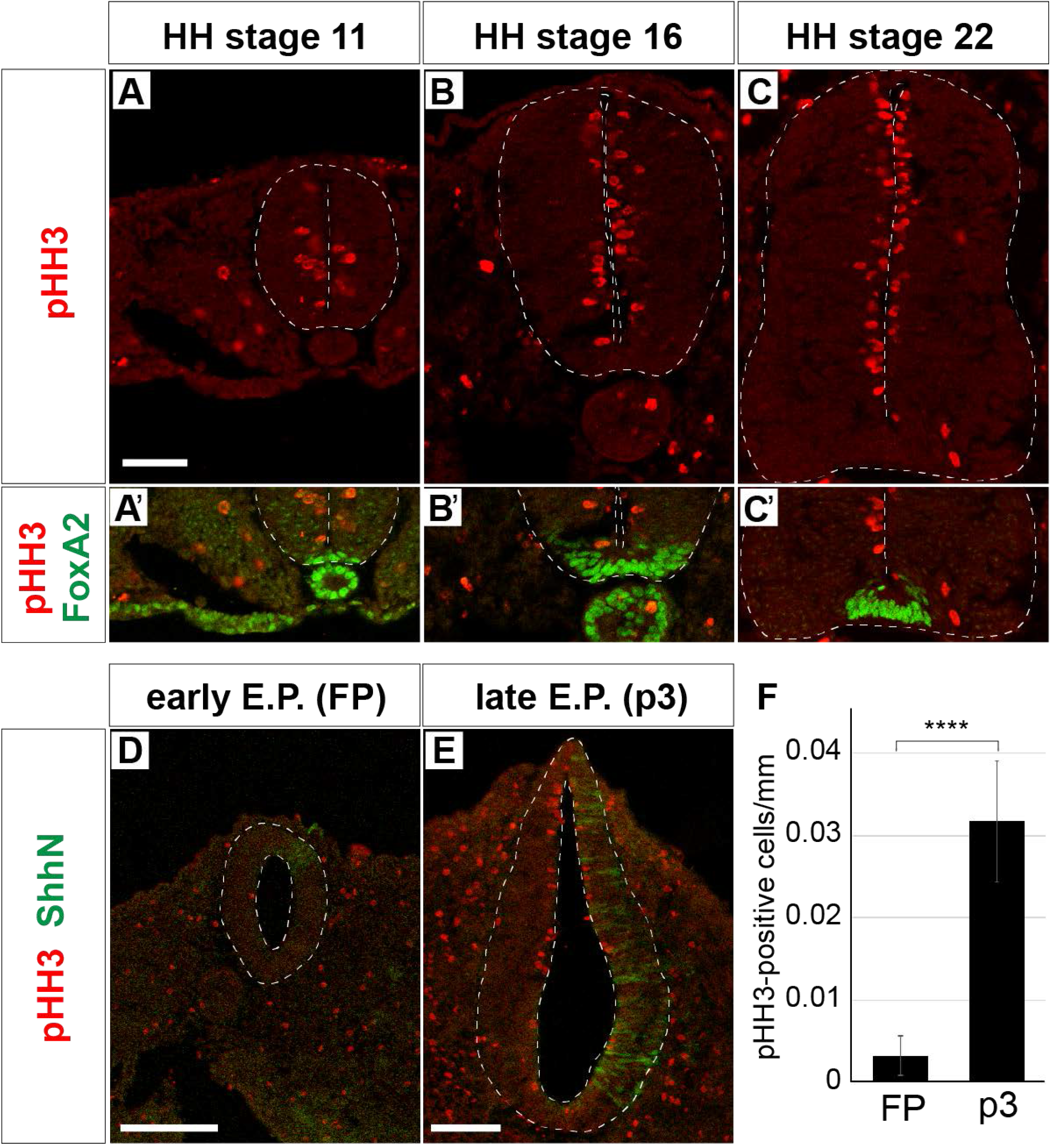
Floor plate cells are not proliferative unlike those of other neural domains. (A-C’) Expression of pHH3 and FoxA2 in neural tube sections at HH stage 11 (A,A’), 16 (B,B’) and 22 (C,C’). pHH3-positive cells were not detected in the FP, where FoxA2 is highly expressed (A’,B’,C’). (D-F) FP cells were less proliferative than p3-interneuron progenitor cells. The neural tube cells differentiate into FP or p3 cells after time-lagged forced expression of ShhN. The expression plasmid for ShhN was electroporated either at HH stage 9 (early E.P.; D) or stage 12 (late E.P.; E) and the embryos were cultured for 48 hours (D) or for 36 hours (E) when they reached HH stage 22 for analysis by immunohistochemistry with the GFP and pHH3 antibodies. Early electroporation (D) led to neural tube differentiation into the FP, whereas late electroporation (E) led to differentiation into p3-interneuron progenitor cells (Ribes et al., 2010; Sasai et al., 2014). (F) Quantitative data for (D) and (E). The positive cells for pHH3 were counted, and the positive rate along the apical surface was presented. Scale bars in (A) for (A-C’),(D),(E) = 50 μm.

Quantitative evaluation of the rate of proliferation of FP cells was difficult because of the small size of the FP domain. We therefore used overexpression analysis to obtain an increased number of FP cells. We previously showed that overexpression of Shh in the chick neural tube at different time points results in a distinct cell fate determination; early electroporation of Shh (i.e. at HH stage 9) leads to the differentiation of the whole neural tube into the FP identity. Moreover, late electroporation (i.e. at HH stage 11) leads to Nkx2.2-positive p3 identity (Ribes et al., 2010; Sasai et al., 2014) in the entire neural tube, providing a good comparison with the FP.

By using this electroporation system, we compared the number of pHH3-positive cells throughout the neural tube sections. The neural tube was significantly smaller in samples undergoing early electroporation than in those undergoing late electroporation of ShhN (Figures 1D-F). In addition, the rate of pHH3-positive apical cells was significantly higher in p3 cells induced by late Shh (Figures 1D-F), suggesting that each domain has a different proliferation rate, and the FP has a low proliferation rate.

Taken together, these results indicate that the cell proliferation activity is significantly lower in FP cells in the neural tube.

### mTOR signal induces the cell proliferation in the neural tube, and is inactive in the FP

We next explored the mechanisms underlying the regulation of cell proliferation in the neural tube. Because the mTOR signaling pathway is important for cell proliferation in many biological contexts (Saxton and Sabatini, 2017), we speculated that the mTOR signal was also involved in regulating the proliferation of neural progenitor cells during neural tube development.

We examined the distribution of the cells active for the mTOR signal along the dorsal-ventral axis of the neural tube. For this purpose, we evaluated two markers of mTOR activity, phospho-p70S6K (p-p70S6K) and its downstream regulator phospho-S6 ribosomal protein (Ser235/236) (hereafter pS6), by immunohistochemisry, at the anterior thoracic level of neural tube of chick (Figures 2A-C,E-G,I-K,M-O,Q-S,U-W,Y-AA) and mouse embryos (Figures 2D,H,L,P,T,X,AB).

**Figure 2.**
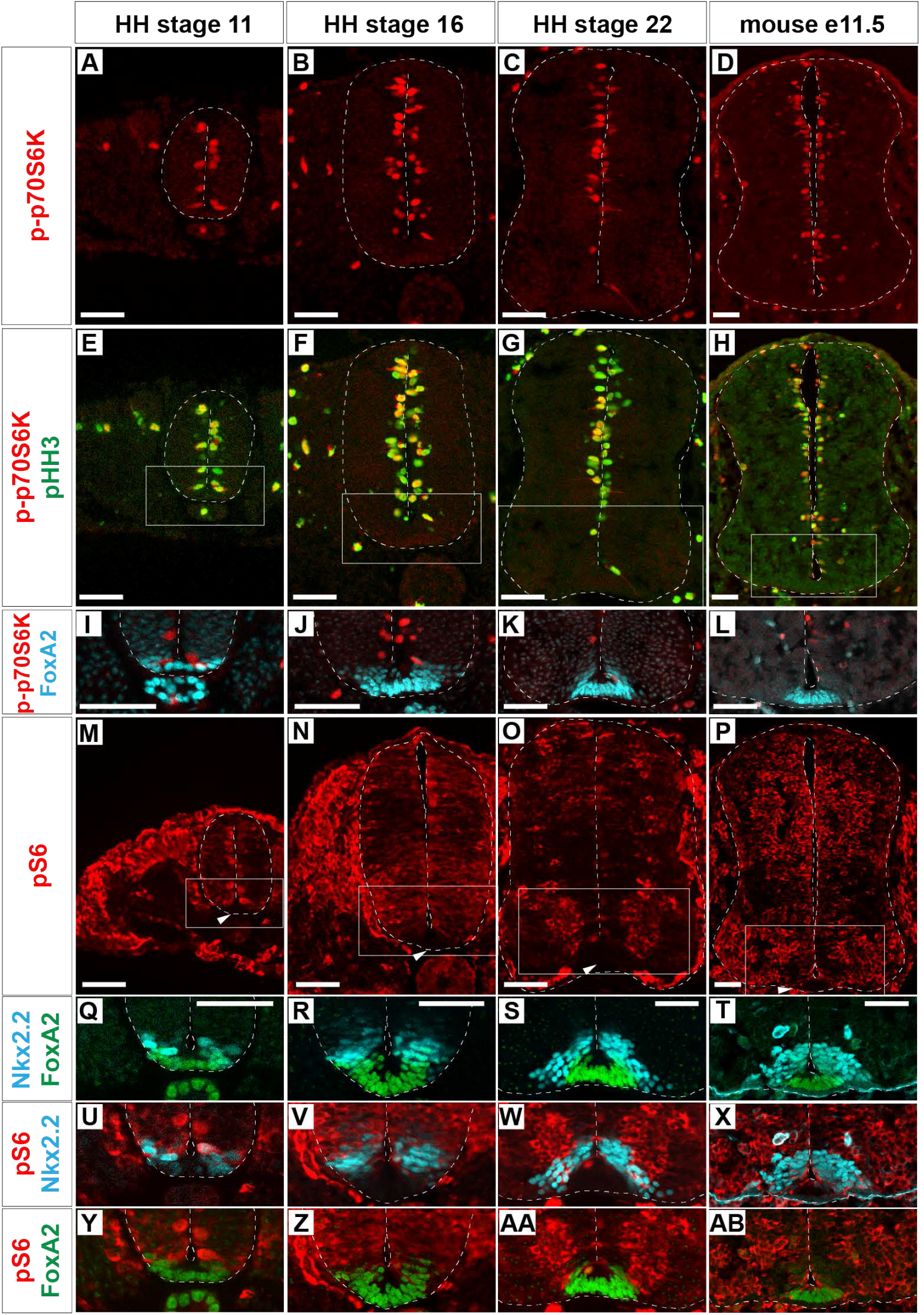
mTOR signal is negative in the floor plate. (A-L) p-p70S6K (A-L), pHH3 (E-H) and FoxA2 (I-L) expression was identified by immunohistochemistry at HH stages 11 (A,E,I), 16 (B,F,J) and 22 (C,G,K) of chick and at e11.5 (D,H,L) mouse neural tube sections. (M-AB) pS6-positive cells (red; M-P,U-AB) were analysed with those of Nkx2.2 (blue; Q-X) and FoxA2 (green; Q-T,Y-AB). (I),(J),(K) and (L) correspond to the areas surrounded by rectangles in (E),(F),(G) and (H), respectively. (Q,U,Y),(R,V,Z),(S,W,AA) and (T,X,AB) correspond to the areas surrounded by rectangles in (M),(N),(O) and (P), respectively. Scale bars = 50 μm. The FP area is indicated by arrowheads (M-P).

p-p70S6K-positive cells were distributed at the apical region of the neural tube at any stages of chick (Figures 2A-C) and mouse (Figure 2D) neural tube. Moreover, importantly, all p-p70S6K-positive cells are included by the pHH3-positive cells (Figures 2E-H), suggesting that the mTOR signal is deeply involved in cell proliferation.

pS6 was detected at the apical domain of the neural tube at HH stage 11 (Figure 2M). At HH stage 16, pS6 was found almost throughout the neural tube with variations in signal intensity (Figure 2N). At HH stage 22, pS6 was detected at the transition zone between progenitor and postmitotic neurons (Figure 2O). On the other hand, in e11.5 mouse neural tube, a strong signal of pS6 was found in the progenitor regions (Figure 2P). Thus the active area for mTOR signal becomes broader at the downstream level as the development progresses, and a species-specific distribution of pS6 was found in the neural tube.

Although mTOR signaling activation, as detected by pS6 expression, was dynamic, neither p-p70S6K nor pS6 were detected in the ventral-most domain at any stage (Figures 2I-L,Q-AB). To more precisely identify pS6-positive cells, pS6-positive domains were compared with FoxA2 and Nkx2.2 expressing domains (Ribes et al., 2010; Sasai et al., 2014). This analysis was performed considering that FoxA2 is weakly expressed in the Nkx2.2-positive p3 domain (Figures 2Q-T), and the *bona fide* FP region is defined by FoxA2-positive and Nkx2.2-negative regions (Ribes et al., 2010; Sasai et al., 2014). The results showed that the ventral end of pS6 expression coincided with the p3 domain, suggesting that the mTOR signal is active in almost all domains in the progenitor regions of the neural tube, but not in the FP (Figures 2U-AB).

Taken together, these results indicated that mTOR signaling is involved in the proliferation of neural progenitor cells, but not FP cells, during neural tube development.

### FoxA2 blocks cell proliferation through negative regulation of mTOR signal

We next focused on the function of FoxA2, a forkhead-type transcription factor, in the regulation of cell proliferation and mTOR signalling in the FP (Ang et al., 1993; Sasaki and Hogan, 1994). FoxA2, which is dominantly expressed in the FP, is one of the primary responsive genes of Shh (Kutejova et al., 2016; Vokes et al., 2007) and is essential for FP differentiation (Placzek and Briscoe, 2005; Sasaki and Hogan, 1994). To understand the involvement of FoxA2 in cell proliferation and mTOR signalling, FoxA2 was overexpressed at HH stage 11 on one side of the neural tube, and the phenotypes were analysed at 48 hpt. We found the FoxA2-overexpressing side was significantly smaller than the control side (Figures 3A-B’). Consistently, the number of pHH3-positive cells was significantly lower in FoxA2-overexpressing cells than in the control GFP-expressing neural tube, suggesting that FoxA2 blocks cell cycle progression (Figures 3A-B’). The cell positive for p-p70S6K (Figures 3D-E’) and pS6 (Figures 3G-H’) were also fewer in the FoxA2-overexpressing side, suggesting that mTOR signaling was inactivated by FoxA2. Conversely, co-expression of CA-mTOR with FoxA2 restored cell proliferation, as characterized by pHH3 expression compared with that in cells expressing FoxA2 alone (Figures 3B-C’). The mTOR signal was also partly recovered in the co-electroporated neural tube (Figures 3F,F’,I,I’). These results suggest that the negative effect of FoxA2 on mTOR signalling is rescued by CA-mTOR, and FoxA2 resides upstream of mTOR.

**Figure 3.**
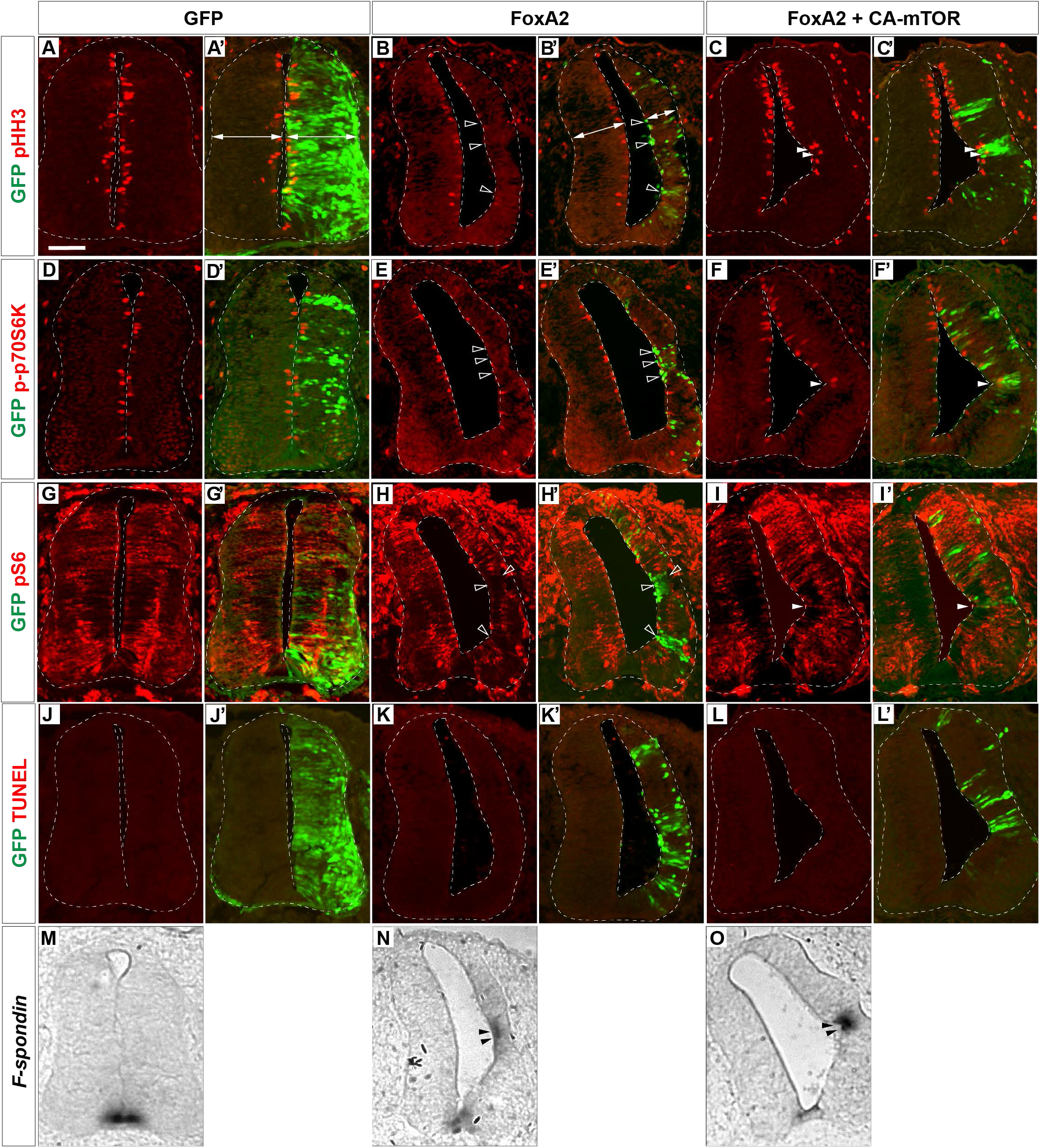
FoxA2 negatively regulates the cell proliferation by blocking the mTOR signal. FoxA2 blocks phosphorylation of p70S6K and S6, and proliferation of the cells without inducing programmed cell death. Plasmids expressing control GFP (A,A’,D,D’,G,G’,J,J’,M), FoxA2 (B,B’,E,E’,H,H’,K,K’,N) of FoxA2 together with CA-mTOR (C,C’,F,F’,I,I’,L,L’,O) were electroporated into one side of the neural tube of HH stage 12 embryos and the phenotypes were analysed at 48 hpt by immunohistochemistry with pHH3 (A-C’), p-p70S6K (D-F’), pS6 (G-I’) and GFP (A’,B’,C’,D’,E’,F’,G’,H’,I’,J’,K’,L’) antibodies, or by a TUNEL assay (J-L’). The merged cells of pHH3 (C,C’), p-70S6K (F,F’) or pS6 (I,I’) with GFP expression are indicated by filled arrowheads, and the pHH3- (B,B’), p-p70S6K- (E,E’) and pS6- (H,H’) negative on GFP-positive cells are indicated by outlined arrowheads. The medio-lateral distances are indicated by double arrows. (M-O) The cell fate determination for FP by FoxA2 is not altered by CA-mTOR. *F-spondin*-positive cells were identified by *in situ* hybridisation. The *F-spondin* expression ectopically induced by FoxA2 is indicated by filled arrowheads (N,O). Scale bar = 50 μm.

We asked if the changes of cell number were mediated by apoptosis, and performed a terminal deoxynucleotidyl transferase dUTP nick end labeling (TUNEL) assay. However, no increasing positive signals were detected in any electroporation (Figures 3J-L’), suggesting that programmed cell death was not the main cause of the alterations in cell numbers.

We finally asked if the cell fate change was involved in the mTOR signalling, and carried out an in situ hybridisation with the FP gene *F-spondin* (Burstyn-Cohen et al., 1999; Klar et al., 1992) probe. As a result, we found the ectopic *F-spondin* expression in both neural tubes electroporated with sole FoxA2 (Figure 3N) or coelectroporation of FoxA2 and CA-mTOR (Figure 3O), whereas the endogenous

Taken together, these results indicate that FoxA2 negatively regulates cell proliferation by blocking the mTOR signal upstream of mTOR.

### RNF152 is expressed in the FP and is a target gene of FoxA2

The role of FoxA2 as a transcription factor led us to hypothesize that FoxA2 induces the expression of gene(s) that directly and negatively regulate mTOR signalling and cell proliferation. To identify such negative regulators of mTOR signalling expressed in the FP, we performed reverse transcription quantitative PCR (RT-qPCR) screening in chick neural explants.

Neural explants treated with a high concentration of Shh (hereafter denoted as Shh^H^; see Materials and Methods for the definition of “high concentration”) differentiate into the FP, whereas explants exposed to a low concentration of Shh (Shh^L^) tend to differentiate into motor neurons and V3 interneurons (Dessaud et al., 2010; Ribes et al., 2010; Sasai et al., 2014). RNA was extracted from explants exposed to Shh^L^ or Shh^H^ for 48 h, and gene expression levels were compared with those of explants without Shh by qPCR focusing on the components of mTORC1 (Laplante and Sabatini, 2009, 2012, 2013) (see Supplementary Table 1 for primer sequences). The results showed that expression of most of the genes was not affected by the presence or absence of Shh (Figure 4A). However, *RNF152*, which encodes an E3 ubiquitin ligase (Deng et al., 2019; Deng et al., 2015), was strongly induced in Shh^H^ explants with a weaker induction with Shh^L^, suggesting that *RNF152* was expressed preferentially in the FP.

**Figure 4.**
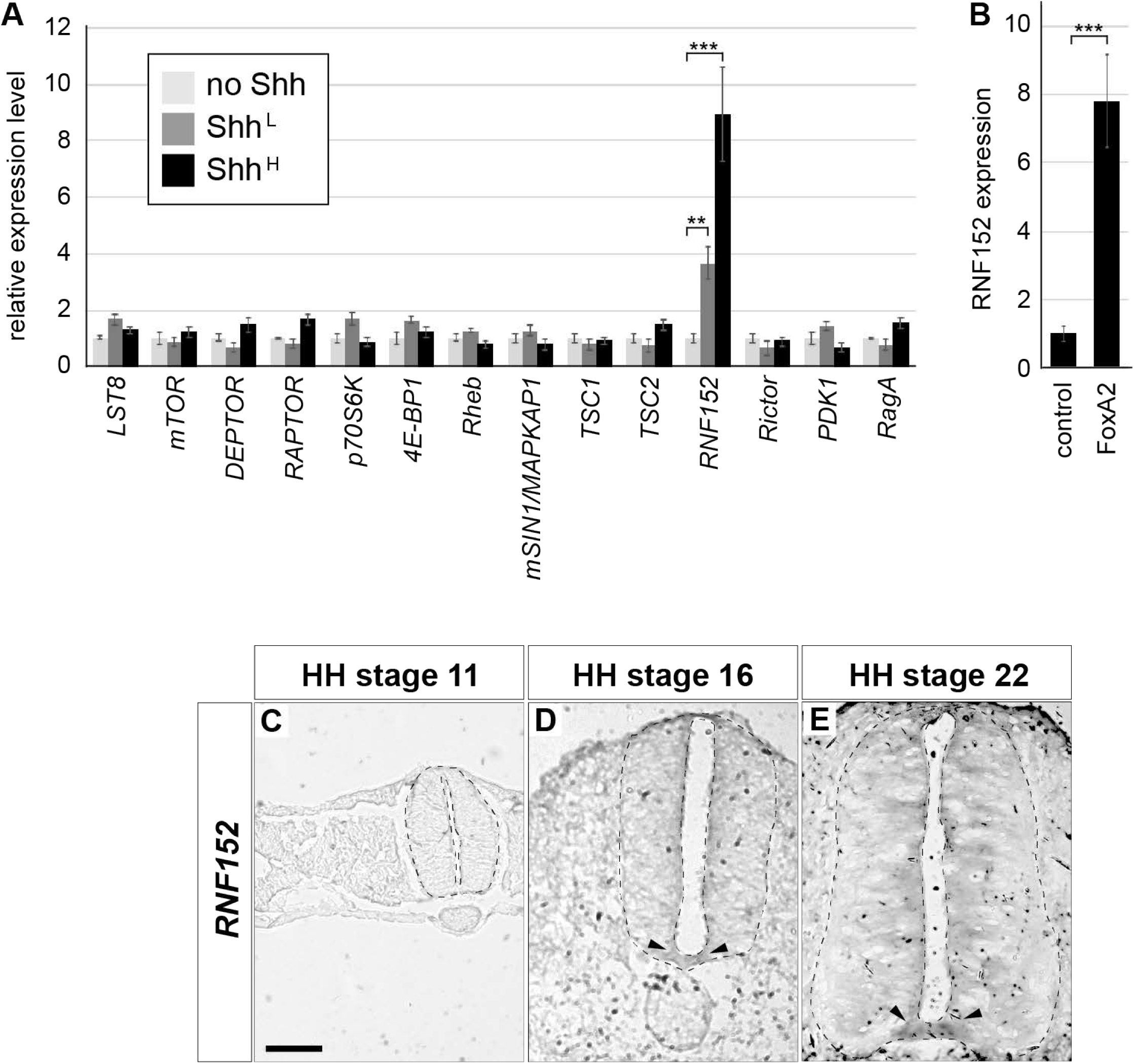
*RNF152* is one of target genes of FoxA2, and is expressed in the FP. (A) *RNF152* is a responsive gene for Shh. RT-qPCR analysis of genes related to the mTOR signal. Chick neural explants treated with control medium or in the presence of Shh^L^ or Shh^H^ for 48 hours were analysed using the indicated gene primers. (B) *RNF152* is a target gene of FoxA2. Explants electroporated with FoxA2 were cultured for 48 hours and the expression of *RNF152* was analysed by RT-qPCR. (C-E) *RNF152* is expressed in the FP. Sections of the neural tube were analysed by *in situ* hybridisation with the *RNF152* probe at HH stage 11 (C), 16 (D) and 22 (E). The FP expression is indicated by arrowheads (D,E). Scale bar = 50 μm.

The regulatory region of the *RNF152* gene contains a FoxA2 binding region (Metzakopian et al., 2012). We therefore prepared explants electroporated with *FoxA2*, and compared gene expression with that of control-GFP electroporated explants by RT-qPCR (Figure 4B). The *RNF152* transcription level was significantly higher in FoxA2-overexpressing explants than in GFP-electroporated explants, suggesting that *RNF152* is a target gene of FoxA2.

To identify the spatial expression of *RNF152* in the neural tube, we performed an *in situ* hybridization analysis. Although the *RNF152* expression was not detected at HH stage 11 (Figure 4C), the expression was found in the FP at HH stages 16 and 22 with a lower level of expression in the apical region of the neural tube (Figures 4D,E).

Taken together, these findings identified *RNF152* as potential negative regulator of the mTOR signal.

### RNF152 negatively regulates cell proliferation through the mTOR signalling pathway

We next attempted to analyse the function of RNF152 in the cell proliferation of the neural tube. Because *RNF152* is expressed in the FP, we first investigated whether RNF152 is involved in FP differentiation. We electroporated the expression plasmid for RNF152 and analysed the FoxA2 expression by immunohistochemistry. However, the expression was not affected by the RNF152 overexpression, suggesting that RNF152 *per se* is not involved in FP fate determination (Figures 5A,A’).

**Figure 5.**
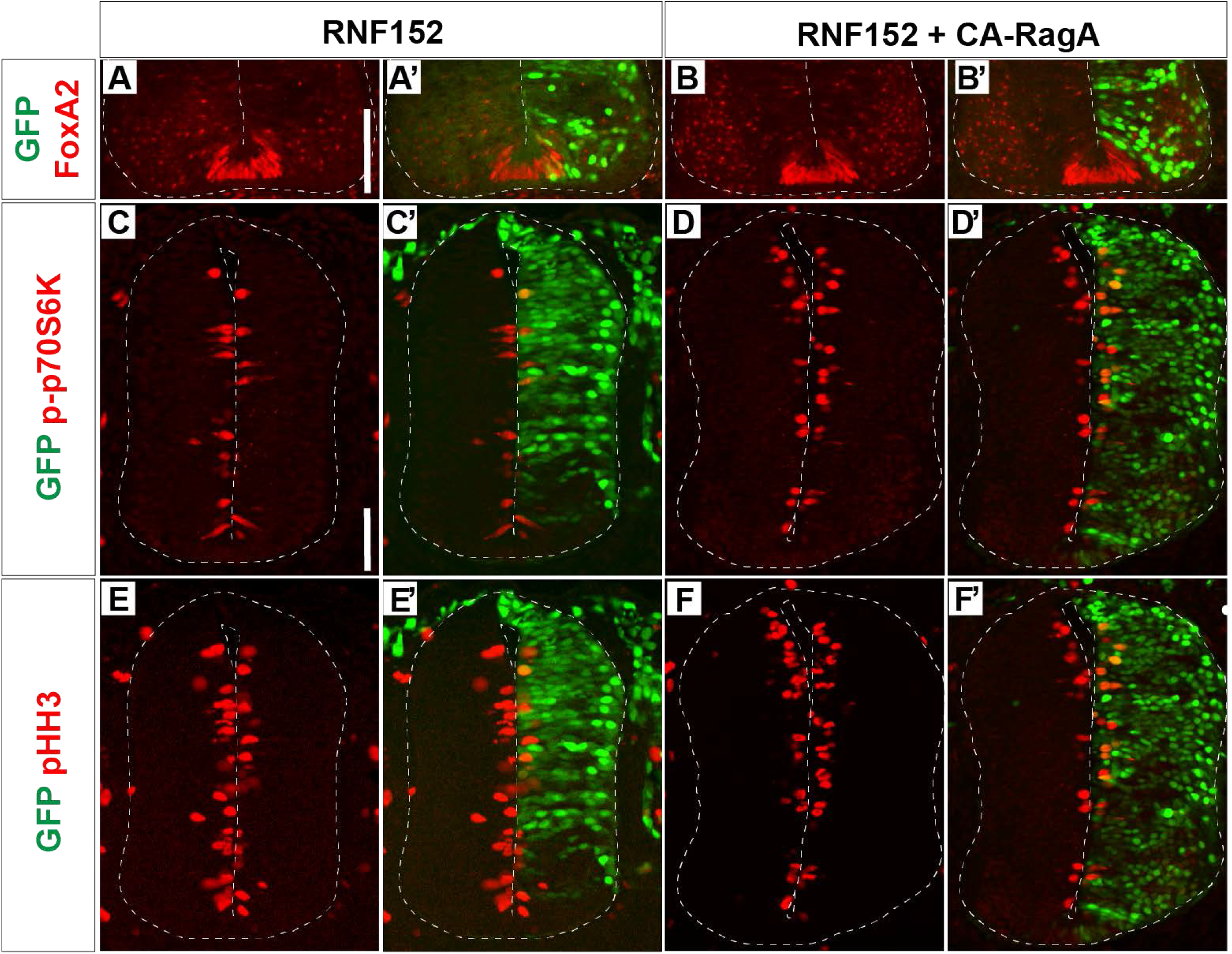
RNF152 negatively regulates the cell proliferation through the mTOR signalling pathway. (A-F) RNF152 negatively regulates mTOR signalling and cell proliferation without altering the cell fate of the FP. The expression plasmids carrying RNF152 (A,A’,C,C’) or RNF152 together with CA-RagA (B,B’,D,D’) were electroporated at HH stage 12 and the FoxA2 (A-B’), p-p70S6K (C-D’), pHH3 (E-F’), and GFP (A’,B’,C’,D’,E’,F’) expression was analysed by immunohistochemistry at 48 hpt. (E-H’) Cell proliferation is regulated by activation of RagA. DN-RagA (E,E’,G,G’) or CA-RagA (F,F’,H,H’) was electroporated at HH stage 12 and phenotypes were analysed at 48 hpt with pHH3 (E-F’), FoxA2 (G-H’) and GFP (E’,F’,G’,H’) antibodies. Scale bars in (A) for (A-B’) and in (C) for (C-F’) = 50 μm.

The *RNF152* gene encodes an E3 ubiquitin ligase targeting the small GTPase RagA (Deng et al., 2019; Deng et al., 2015; Kim et al., 2008), and the GTP-bound active form of RagA positively regulates the mTOR signalling pathway (Efeyan et al., 2014; Shaw, 2008). Actually the expression of the dominant-negative RagA (DN-RagA) blocks cell proliferation in the electroporated cells while the constitutively-active RagA (CA-RagA) activates it, without changing the FP cell fate (Supplementary Figure 2). RNF152 was therefore expected to act as a negative regulator of the mTOR signalling pathway by blocking RagA activity. To prove this hypothesis, p-p70S6K expression was analysed in cells overexpressing RNF152. The results showed p-p70S6K was downregulated in response to RNF152 overexpression (Figure 5C,C’), suggesting that RNF152 is a negative regulator of mTOR signalling. We further asked if the effect of RNF152 on cell proliferation in the neural tube, and analysed the expression of pHH3 by immunohistochemistry, which showed that the number of pHH3-positive cells was significantly lower in RNF152-overexpressed side than in the unelectroporated side (Figures 5E,E’). Therefore, RNF152 negatively regulates cell proliferation via blocking the mTOR signalling pathway.

We next asked if the effect of RNF152 can be rescued by hyperactivation of RagA. We therefore electroporated CA-RagA together with RNF152, and investigated the expression of FoxA2, p-p70S6K and pHH3. As a result, the number of p-p70S6K- and pHH3-positive cells was significantly higher in cells co-electroporated with CA-RagA and RNF152 than in those overexpressing RNF152 alone (Figure 5D,D’,F,F’), while FoxA2 expression was unchanged (Figure 5B,B’), suggesting that RagA resides downstream of RNF152, and controls cell proliferation by antagonizing RNF152.

In summary, RNF152 negatively regulated cell proliferation by blocking mTOR signalling upstream of RagA.

### Blocking RNF152 expression leads to aberrant cell division in the FP

To elucidate the function of RNF152 in mTOR signaling and FP cell proliferation, we designed a loss-of-function experiment to inhibit the RNF152 expression by si-RNA. We electroporated *si-control* or *si-RNF152* in the ventral region of the neural tube together with the GFP-expressing plasmid at HH stage 10, and cultured the embryos for 48 h to reach HH stage 18.

While no ventral expansion of pS6 was found by the *si-control* electroporation (Figure 6A,A’), *si-RNF152* induced aberrant pS6 expression in the FP (Figures 6B,B’), suggesting that the mTOR signal can be reverted by inhibiting RNF152. Moreover, pHH3 was found in the midline cells (Figure 6G), which was never expressed in the *si-control*-electroporated neural tube (Figures 6F,G). This pHH3-positive cells coexpressed FoxA2 (Figure 6G”), suggesting that the aberrant pHH3 expression was induced by the perturbation of RNF152 expression.

**Figure 6.**
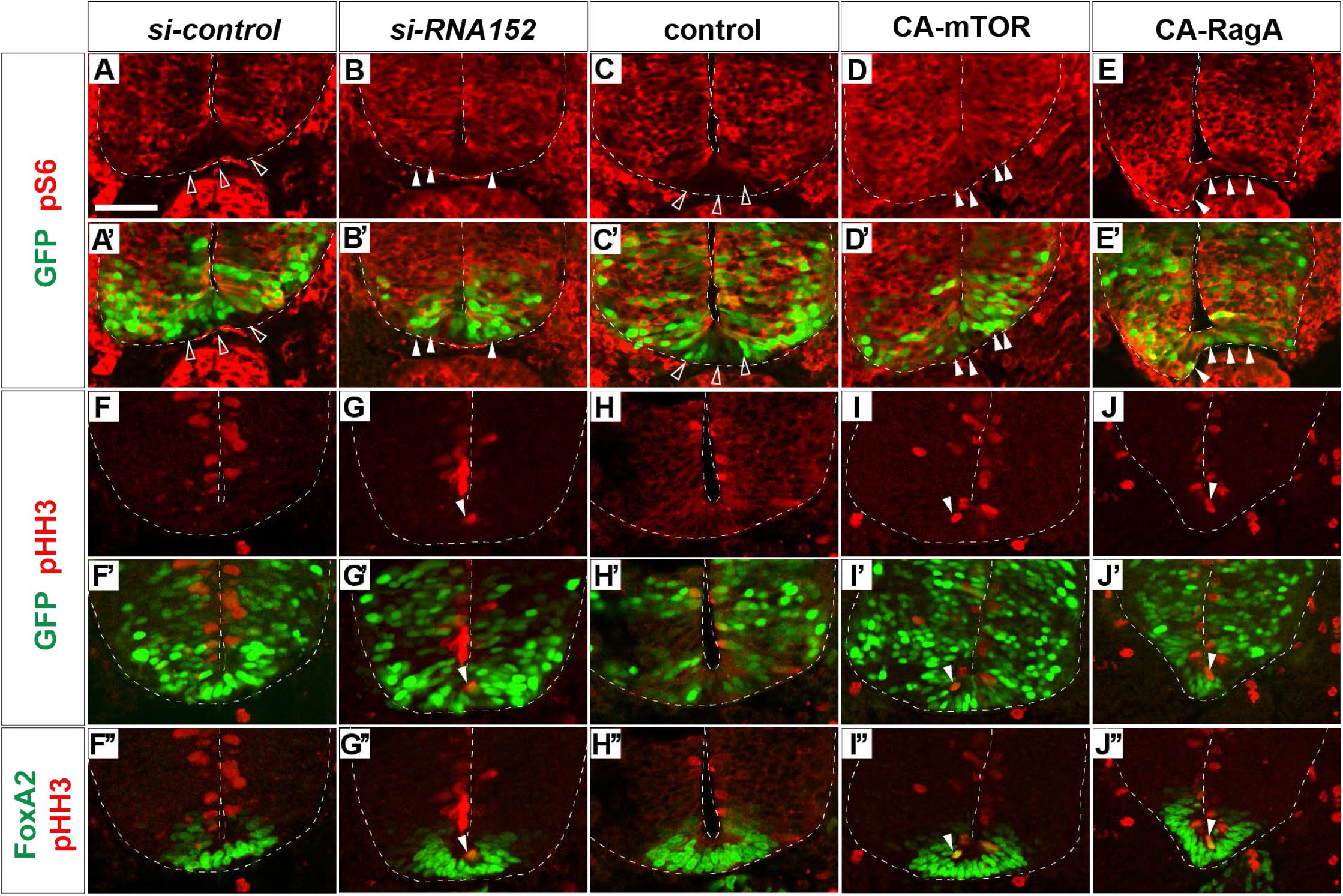
Blocking RNF152 expression leads to aberrant mTOR signal upregulation and cell division in the floor plate. (A-B’,F-G”) Knockdown of RNF152 by *si-RNA* caused aberrant mTOR activation and the appearance of pHH3-positive cells. *si-control* (A,A’,F,F’,F”) or *si-RNF152* (B,B’,G,G’,G”) were electroporated in the FP at HH stage 10 and embryos were analysed at 48 hpt with pS6 (A-B’), pHH3 (F-G”), FoxA2 (F”,G”) and GFP (A’,B’,F’,G’) antibodies. (C-E’,H-J”) Activation of mTOR signal induces aberrant cell division. The plasmids of control (C,C’,H,H’,H”), CA-mTOR (D,D’,I,I’,I”), or CA-RagA (E,E’,J,J’,J”) was electroporated in the same way as in (A,B) and analysed with pS6 (C-E’), pHH3 (H-J”) and FoxA2 (H”,I”,J”) and GFP antibodies (C’,D’,E’,H’,I’,J’). The affected areas are indicated by filled arrowheads and outlined arrowheads. Scale bar = 50 μm.

We confirmed that the activation of mTOR signal induced the ectopic pHH3 expression in the FP region. We electroporated control, CA-mTOR or CA-RagA in the ventral neural tube, and checked the expression of pS6 and pHH3. As expected, the pS6 expression was found in the FP region in CA-mTOR and CA-RagA electroporation while no expansion was found the control electroporation (Figure 6C-E’). Moreover, pHH3 expression, which was not found in the FP upon the electroporation of the control plasmid, was found in the midline cells. Moreover, the pHH3-positive cells coexpressed FoxA2, suggesting that the ectopic pHH3 expression did not change the FP cell fate (Figures 6H”,I”,J”). Finally, the FoxA2 expression domain did not change by the electroporation of CA-mTOR or CA-RagA, suggesting that the FoxA2 expression was regulated at the upstream level of the mTOR signal.

Altogether, RNF152 is essential for inhibiting the cell proliferation, and blocking the function of RNF152 either by *si-RNA* or by activating mTOR signal induced the aberrant cell division in the FP.

## Discussion

### RNF152 is a negative regulator of mTOR signalling in neural tube development

In the present study, we demonstrated that activation of the mTOR signalling pathway promotes cell proliferation in the neural tube. mTOR signalling is inactivated in the FP, which corresponds to the low proliferation rate of FP cells. FoxA2 is an essential transcription factor that restricts cell proliferation, and this negative regulation is mediated by RNF152, an E3 ubiquitin ligase that targets the mTOR pathway component RagA and a target gene of FoxA2.

Although Shh regulates not only pattern formation in the neural tube, but also cell proliferation and tissue growth, FP cells exposed to the highest level of Shh have a low proliferation rate (Kicheva et al., 2014). The present study elucidated the mechanism underlying this regulatory function.

RNF152, a lysosome-anchored E3 ubiquitin ligase (Deng et al., 2019; Zhang et al., 2010) containing RING-finger and transmembrane domains, was initially thought to induce apoptosis (Zhang et al., 2010). Further study showed that RNF152 ubiquitinates and targets the GDP-bound form of RagA for degradation, thereby negatively regulates mTOR signalling (Deng et al., 2015). Consistently, *RNF152* knockout cells exhibit hyperactivation of mTOR signalling (Deng et al., 2015). Moreover, a recent study proposed that RNF152 has an essential function in neurogenesis by regulating *NeuroD* expression (Kumar et al., 2017). Although these findings at the cell level suggest that RNF152 plays essential roles during the entire course of life including embryogenesis, mutant mice devoid of the *RNF152* gene are actually viable (Deng et al., 2015), suggesting the existence of a compensatory mechanism for RNF152 to ensure survival.

On the other hand, RagA, a substrate of RNF152, is essential for embryogenesis; genetic deletion of the *RagA* gene causes morphological and growth defects, and the embryos consequently die at embryonic day 10.5 (Efeyan et al., 2014). This suggests that RagA activation is not exclusively regulated by RNF152, and other factors may be involved in the modulation of RagA activity. To elucidate the critical function of RNF152 in certain aspects of development or at postnatal stages, conditional knockout mice are needed to delete specific functions at a specific space and time.

Consistent with the diverse functions of RNF152 and RagA, the downstream mTOR signal is involved at multiple levels during neural development (Yu and Cui, 2016). In addition to its roles in neurogenesis (Fishwick et al., 2010), mTOR signaling is essential for neural tube closure, as demonstrated in TSC1/2 knockout mice (Kobayashi et al., 2001). Although the critical function of mTOR signalling during neural development is known, an integral understanding of the role of mTOR signalling in this process requires conditional knockout mice to delete its function during each stage of neural development.

### Factors upstream of the mTOR signal

Figure 7 is a schematic of the regulation of cell proliferation in the FP. FoxA2, a target of Shh, induces the expression of downstream target genes including RNF152. RNF152 inactivates the mTOR signalling pathway, thereby negatively regulates cell proliferation. In this sense, our present study linked the two signalling pathway of Shh and mTOR.

**Figure 7.**
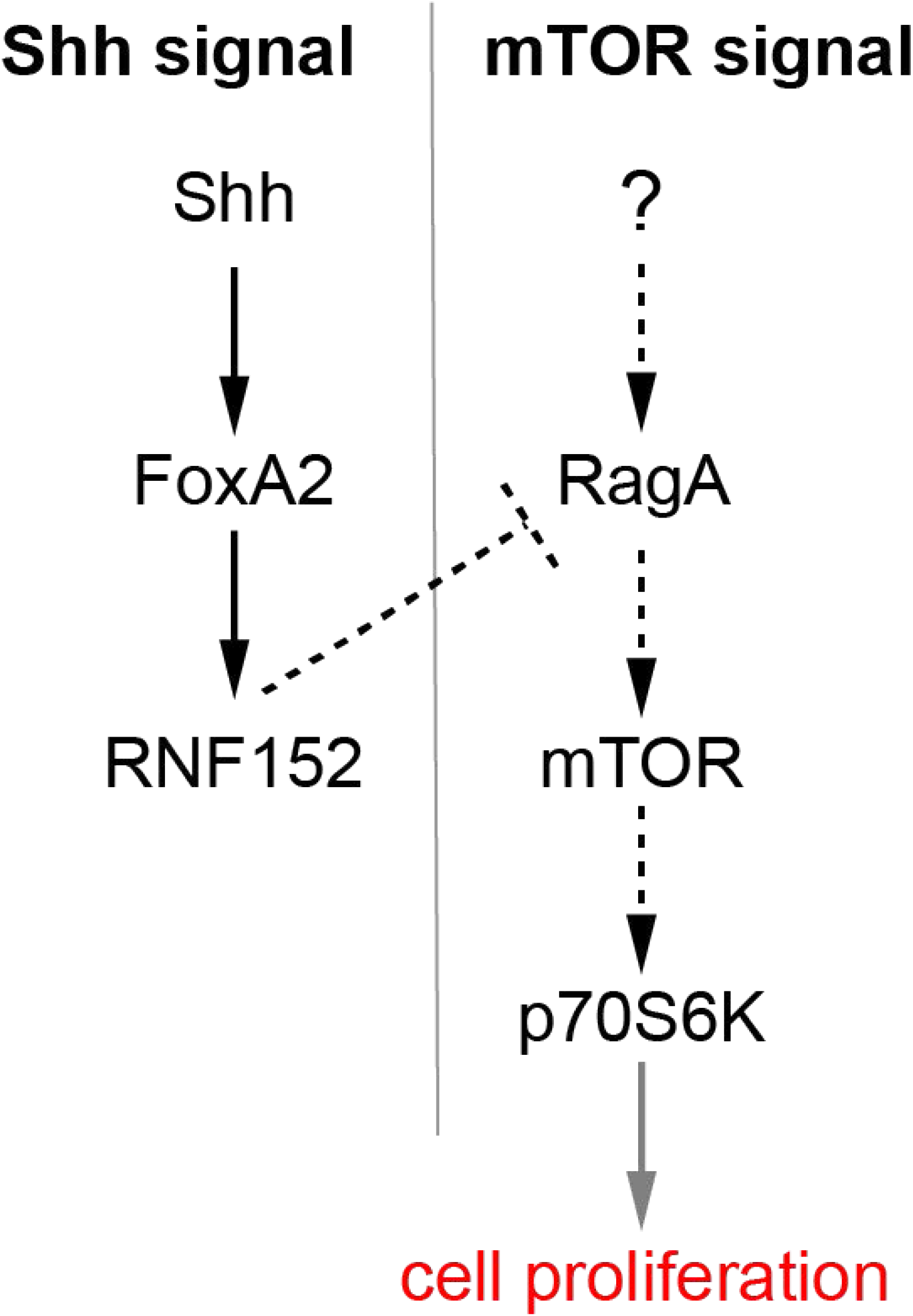
A regulatory loop composed of Shh, FoxA2 and RNF152 modulates FP cell proliferation. *FoxA2* expression is induced by Shh, whereas *RNF152* is a target gene of FoxA2. RNF152 blocks the cell proliferation through by negatively regulating mTOR signalling. Transcriptional regulation is indicated by solid arrows; activation, and inactivation with protein interactions or modifications are indicated by dotted arrows; the regulation of cell proliferation by the activation of p70S6K is apparently indirect, which is indicated by the gray arrow.

The upstream component of the mTOR pathway that is active during neural tube development remains unidentified. mTOR signalling can be activated by insulin-like growth factor (IGF) (Laplante and Sabatini, 2012). However, the expression of IGF and the activation of mTOR do not occur in parallel during neural tube development. IGF1 is not expressed at a detectable level during neural tube development (NS, unpublished observation). Furthermore, IGF2 and IGF1R (IGF1 receptor) are expressed in somites and in the dorsal part of the neutral tube (Fishwick et al., 2010), whereas mTOR is phosphorylated in the ventral neural tube and in the neural crest (Nie et al., 2018). pAKT, which resides upstream of mTOR, is active at the apical domain of the neural tube and along the dorsoventral axis (Supplementary Figures 1A,B), and later in the commissural axons (Supplementary Figure 1C), which does not correspond to the distribution of the downstream molecule pS6 (Figure 2C,O). These results suggest that mTOR signal is activated dynamically and plays multiple roles during neural tube development; the progenitor cell proliferation is encouraged at earlier stages, while the maturation and/or the migration of the neurons at later stages. Moreover, it is quite possible that more than one upstream factors activate the pathway in a context dependent manner.

Future analyses can focus on the behaviour of single cell at different developmental stages to find the new effector(s), which will elucidate the mechanisms by which diverse mTOR functions are exerted.

## Materials and Methods

### Ethical Statements

All animal experiments were carried out in accordance with the national and domestic legislations. All protocols of the experiments on chick and mouse embryos were approved by the animal research review panel of Nara Institute of Science and Technology (approval numbers 1636 and 1810, respectively).

### Electroporation, immunohistochemistry and *in situ* hybridisation

Chicken eggs were purchased from the Yamagishi Farm (Wakayama Prefecture, Japan), and developmental stages were evaluated according to the Hamburger and Hamilton criteria (Hamburger and Hamilton, 1992). Electroporation was performed with the ECM 830 (BTX) electroporator in the neural tube of embryos using pCIG-based expression plasmids, in which gene expression is induced by the chicken beta-actin promoter (Megason and McMahon, 2002). For the ventral electroporation (Figure 6), electrodes was placed on the embryos and under the embryos. *pCIG-CA-mTOR* was generated by modifying the *pcDNA3-FLAG-mTOR-S2215Y* vector purchased from Addgene (# 69013), which was deposited by Dr. David Sabatini (Grabiner et al., 2014). Detailed information of the plasmids and si-RNAs used in this study is provided in Supplementary Table 2. Embryos were incubated in a 38°C incubator for the indicated times at constant humidity.

Embryos were fixed with 4% paraformaldehyde on ice for 2 hours, and then incubated with 15% sucrose/PBS solution overnight. Embryos were embedded in the OCT compound (Sakura) and sectioned at a thickness of 12 μm (Sakura Finetek, Japan).

Immunohistochemistry and *in situ* hybridization were performed as described previously (Sasai et al., 2014). The antibodies used in this study are listed in Supplementary Table 2.

Timed pregnant mice were purchased from Japan SLC (Shizuoka Prefecture, Japan). Embryos were extracted and processed as described for chick embryos.

### Explants, RNA extraction and RT-qPCR

Intermediate neural explants comprise the uniform type of neural progenitor cells, which are sensitive to patterning factors and are a useful experimental model to recapitulate *in vivo* neural development (Dessaud et al., 2010; Sasai et al., 2014). For preparation, chick embryos were extracted from eggs at HH stage 9, and the intermediate region of the neural plate at the preneural tube level (Delfino-Machin et al., 2005) was excised. If necessary, expression plasmids were overexpressed before extracting the embryos (Figure 4B). Explants were embedded in a pH-adjusted collagen gel with DMEM. The culture medium consisted of DMEM/F-12 (Thermo Fisher Scientific), Mito+Serum Extender (Sigma), and penicillin/streptomycin/glutamine (Wako). Recombinant Shh was prepared in house (Kutejova et al., 2016; Sasai et al., 2014). Shh^H^ was defined as the concentration at which the explants produced a dominant population of Nkx2.2-positive cells with a small subset of Olig2 cells at 24 h. SHH^L^ was defined as 1/4 of the concentration of Shh^H^, producing >70% Olig2-positive cells and a lower number of Nkx2.2-positive cells (Dessaud et al., 2010; Dessaud et al., 2007). At the late 48 h time point, Shh^H^ explants differentiated into the FP, whereas Shh^L^ induced motor neuron differentiation, as characterized by Islet1 expression (Ribes et al., 2010; Yatsuzuka, 2018).

For RT-qPCR, RNA was extracted using the NucleoSpin RNA extraction kit (Macherey-Nagel U0955), and cDNA was synthesized using the PrimeScript II cDNA synthesis kit (TaKaRa 6210). The qPCR reaction mixtures were prepared with SYBR FAST qPCR master mix (KAPA KR0389), and PCR amplification was quantified by LightCycler 96 (Roche).

### Images collection and statistical analysis

The immunofluorescent and *in situ* hybridisation images were captured with the LSM 710 confocal microscope and axiocam digital camera (Carl Zeiss), and were processed by Photoshop CC (Adobe) and figures were integrated by Illustrator CC (Adobe). Statistical analyses were carried out by using Prism (GraphPad). Statistical data are presented as mean values ± s.e.m., and significance (**; p<0.01, ***; p<0.001, ****; p<0.0001 or n.s.; not significant. Statistical analyses between two groups were carried out with two-tailed t-test.

## Data accessibility

All the data are available in the main text, figures and the supplementary materials.

## Author contributions

NS conceived the project. MK and NS performed experiments, and analysed the data. NS wrote the manuscript.

## Acknowledgements

The authors thank DSHB (Developmental Studies Hybridoma Bank) at the University of Iowa, USA, and Addgene (the non-profit plasmid repository) for materials, Michinori Toriyama and the laboratory members for support and discussions.

## Supplementary Information

**Supplementary Figure 1.**
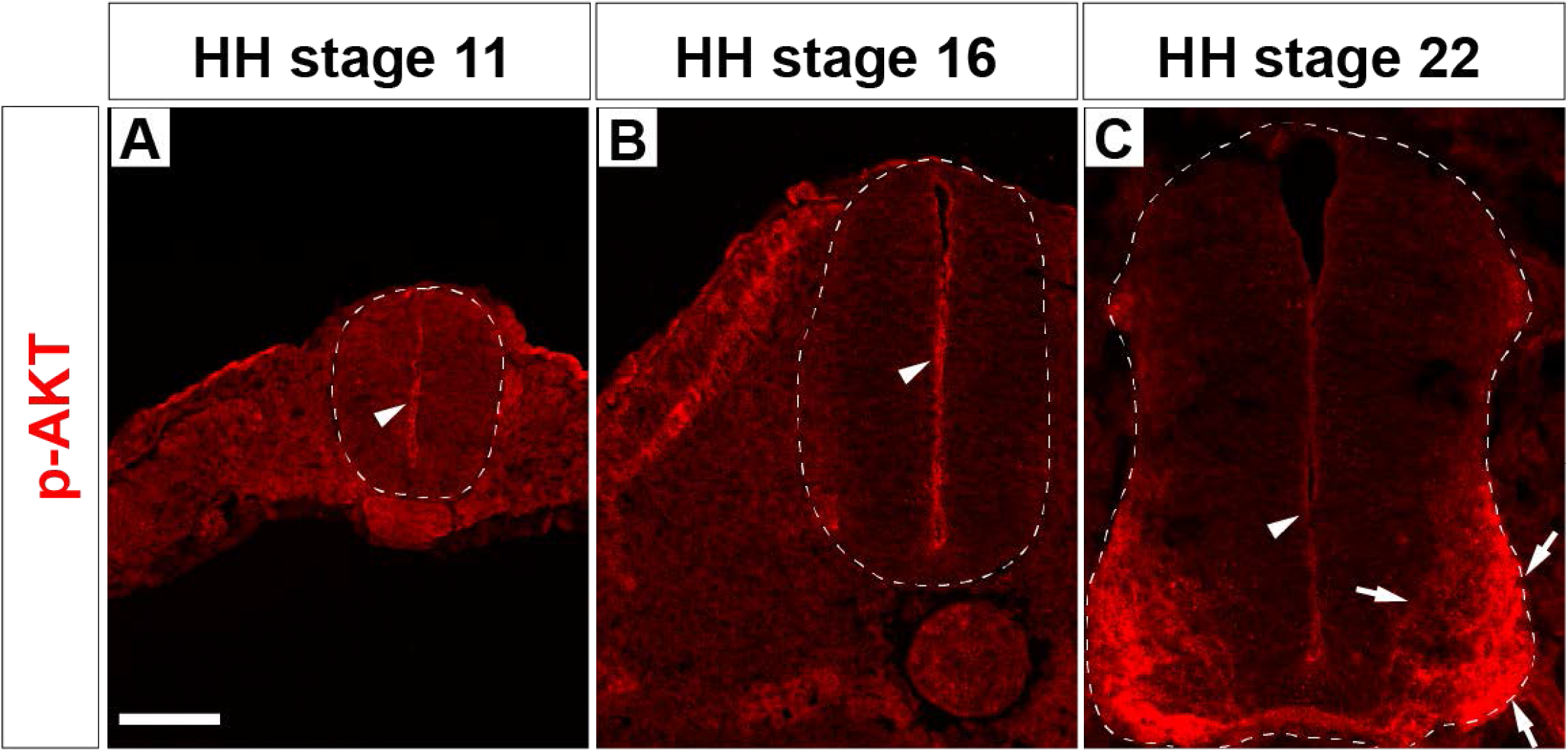
pAKT is localised at the apical domain and at commissural neurons. pAKT-positive cells are identified by immunohistochemistry in chick neural tube sections HH stages 11 (A), 16 (B) and 22 (C). Expression in the apical domain and in the commissural axons are indicated by arrowheads and arrows, respectively. Scale bar = 50 μm.

**Supplementary Figure 2.**
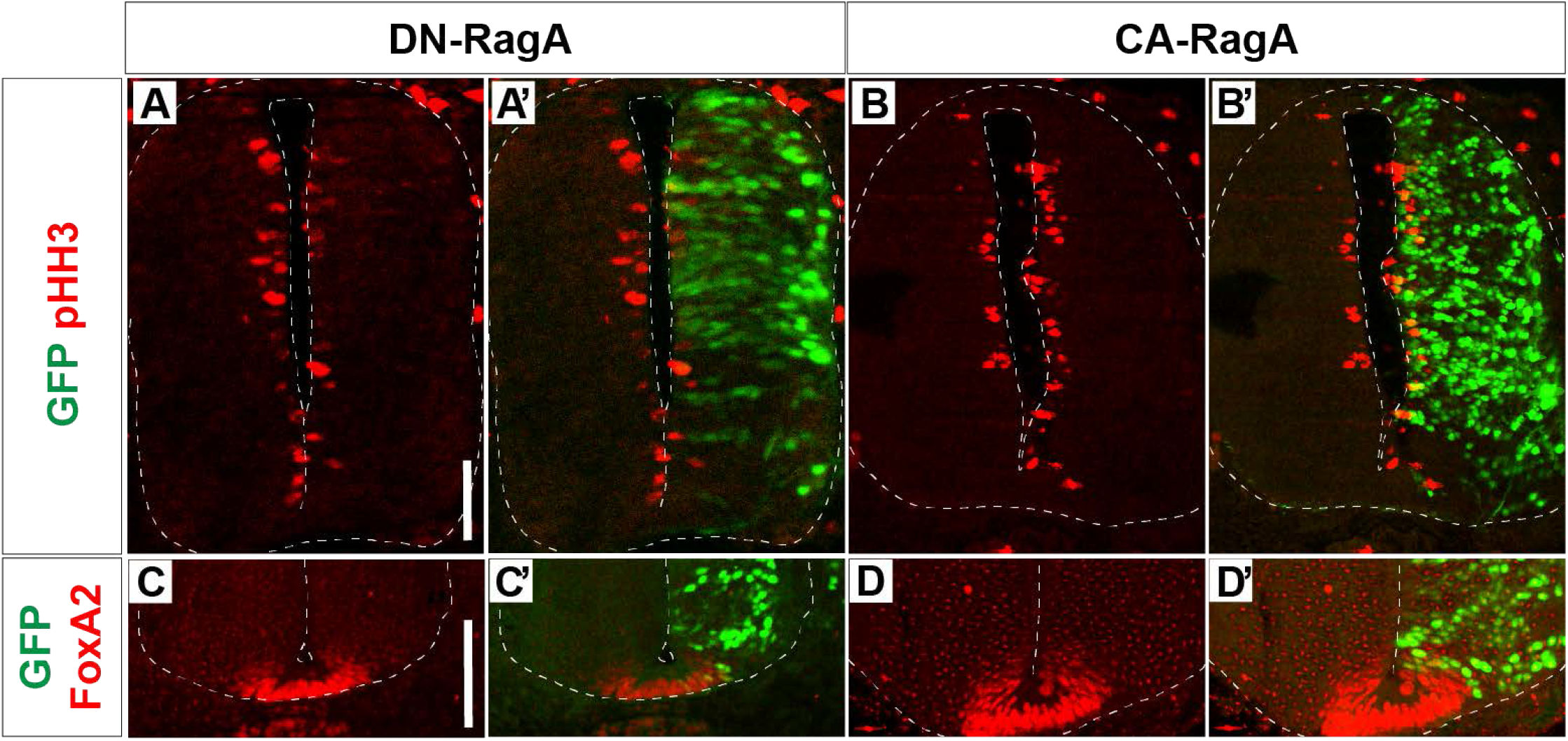
Cell proliferation is regulated by activation of RagA. DN-RagA (A,A’,C,C’) or CA-RagA (B,B’,D,D’) was electroporated at HH stage 12 and phenotypes were analysed at 48 hpt with pHH3 (A-B’), FoxA2 (C-D’) and GFP (A’,B’,C’,D’) antibodies. Scale bars in (A) for (A-B’) and in (C) for (C-D’) = 50 μm.

Supplementary Table 1 Primers for quantitative PCR

Supplementary Table 2 Plasmids, siRNAs and antibodies used in this study

(references)

(Grabiner et al., 2014; Kim et al., 2008; Li et al., 2004; Sasai et al., 2014; Sato et al., 2008; Tabancay et al., 2003; Yatsuzuka, 2018)

